# Multiple mechanisms for overcoming lethal over-initiation of DNA replication

**DOI:** 10.1101/2021.05.06.442943

**Authors:** Mary E. Anderson, Janet L. Smith, Alan D. Grossman

## Abstract

DNA replication is a highly regulated process that is primarily controlled at the step of initiation. In the gram-positive bacterium *Bacillus subtilis* the replication initiator DnaA, is regulated by YabA, which inhibits cooperative binding at the origin. Mutants lacking YabA have increased and asynchronous initiation. We found that under conditions of rapid growth, the *dnaA1* mutation that causes replication over-initiation, was synthetic lethal with a deletion of *yabA*. We isolated several classes of suppressors of the lethal phenotype of the Δ*yabA dnaA1* double mutant. Some suppressors (*dnaC, cshA*) caused a decrease in replication initiation. Others (*relA, nrdR*) stimulate replication elongation. One class of suppressors decreased levels of the replicative helicase, DnaC, thereby limiting replication initiation. We found that decreased levels of helicase were sufficient to decrease replication initiation under fast growth conditions. Our results highlight the multiple mechanisms cells use to regulate DNA replication.

## Introduction

DNA replication is a highly regulated, essential process across all domains of life. All organisms need to control DNA replication based on environmental cues and growth rate to ensure each daughter cell has a complete copy of the genome. Multiple mechanisms are used to coordinate DNA replication with other cellular processes such as metabolism and cell division, as well as to sense external cues that can impact DNA replication. Failure to properly regulate DNA replication can result in a variety of consequences, such as cell division defects, anucleate daughter cells, DNA damage, and in higher organisms, disease (O’Donnell, Langston, and Stillman, 2013; Magdalou *et al*., 2014).

Bacteria typically have a single circular chromosome and initiate bi-directional DNA replication from an origin, finishing roughly 180° opposite, at the terminus. Under favorable, nutrient rich growth conditions, certain bacteria, including *Escherichia coli* and *Bacillus subtilis*, can undergo multifork replication; that is, they initiate a new round of DNA replication before the previous round has finished. Multifork replication results in each daughter cell receiving a chromosome with active replication forks and multiple origins, enabling cells to divide more quickly, while still ensuring each daughter cell receives a completed chromosome (reviewed in Skarstad and Katayama, 2013). Under slow growth conditions bacteria restrict replication initiation. Overinitiation can lead to replication fork collapse and the DNA damage response, as in addition to problems with cell division and chromosome segregation (Katayama, 2001; Bach and Skarstad, 2004; O’Donnell, Langston, and Stillman, 2013; Magdalou *et al*., 2014). This flexibility in regulating the rate of replication initiation gives bacteria the advantageous ability to reliably adjust growth and division based on environmental and internal cues.

In bacteria, DNA replication initiation is primarily regulated by controlling the levels and activity of the replication initiator, DnaA. DnaA is AAA+ ATPase which binds both ADP and ATP (reviewed in Davey *et al*., 2002). The ATP-bound form is active for replication initiation and binds cooperatively to DnaA boxes located near the origin of replication (*oriC*). This cooperative binding causes formation of a nucleoprotein helical filament that melts the AT-rich DNA unwinding element (DUE) (reviewed in Leonard and Grimwade, 2005). In *B. subtilis*, DnaA recruits additional proteins required for chromosome organization and helicase loading (DnaD and DnaB) before the helicase loader (DnaI). These proteins load the hexameric replicative helicase (DnaC) monomer by monomer around the DNA before the remaining replication machinery is recruited to the origin, and DNA replication proceeds bi-directionally (reviewed in Kaguni, 2006; Mott and Berger, 2007; Leonard and Grimwade, 2011).

The protein levels and activity of DnaA directly affect initiation. For example, increased expression of *dnaA* causes overinitiation (Atlung *et al*., 1987; Skarstad *et al*., 1989) and various *dnaA* mutations have been characterized that either enhance or inhibit DNA replication initiation (Moriya *et al*., 1990; Guo *et al*., 1999; Murray and Errington, 2008; Scholefield and Murray, 2013). One such mutation, *dnaA1*, causes a serine to phenylalanine change at amino acid 401F (S401F) causes a temperature sensitive phenotype. Replication initiation is inhibited at non-permissive temperatures where the DnaA1 mutant protein is unstable (Moriya *et al*., 1990).

DnaA is also regulated by mechanisms that prevent its cooperative binding at the origin. In *Escherichia coli*, this is primarily accomplished by regulating the availability of ATP-bound DnaA, either through sequestration or titration (*seqA/datA*) or regulated inactivation of DnaA (RIDA) by Hda (reviewed in Kaguni, 2006; Leonard and Grimwade, 2011). However, in gram-positive bacteria like *B. subtilis*, DnaA is regulated by direct interactions with several proteins that affect cooperative binding, including YabA (Wagner, Marquis, and Rudner, 2009; Merrikh and Grossman, 2011; Scholefield, Errington, and Murray, 2012; Bonilla and Grossman, 2012; Scholefield and Murray, 2013). YabA binds directly to DnaA and prevents the necessary cooperative binding at DnaA boxes near the origin, thereby inhibiting formation of the nucleoprotein filament and melting of the origin (Merrikh and Grossman, 2011; Scholefield and Murray, 2013). YabA also stimulates dissociation of DnaA from *oriC* (Schenk, *et al*., 2017). A null mutation in *yabA* leads to increased and asynchronous DNA replication (Hayashi *et al*., 2005; Noirot-Gros *et al*., 2002; Goranov *et al*., 2009).

We found that the *dnaA1* mutation causes overinitiation at permissive temperature, and that combining *dnaA1* with a null mutation in *yabA* results in a more extreme overinitiation phenotype that was lethal when cells were grown in rich medium. We leveraged this conditional synthetic lethal phenotype to isolate suppressors that restored viability to the cells that had extreme overinitiation of replication, with the aim of elucidating novel mechanisms of DNA replication regulation. By selecting for survival under conditions of rapid growth (LB medium), we identified mutations affecting five different genes that suppressed the lethal phenotype of the Δ*yabA dnaA1* double mutant. We found that null mutations in *cshA* suppressed lethality by decreasing replication initiation, whereas a null mutation in *nrdR* and a mutation affecting the (p)ppGpp synthetase domain of *relA* suppressed lethality by stimulating replication elongation to keep pace with increased initiation. We found that mutations that decrease levels of the replicative helicase, DnaC, were sufficient to limit DNA replication initiation under high initiation conditions, either in the overinitiating Δ*yabA dnaA1* background or under fast growth conditions. In addition to elucidating novel genes that can regulate replication initiation, the results of this screen highlight the multiple ways cells can regulate DNA replication, by employing changes in other cellular processes to compensate for a detrimental overinitiation phenotype.

## Results

### Mutations in *yabA* and *dnaA* cause a synthetic lethal phenotype in rich medium

*dnaA1*(S401F) causes a temperature sensitive phenotype that results in a loss of replication initiation at non-permissive temperatures (Moriya *et al*., 1990). DnaA is also a transcription factor and at permissive temperature, the DnaA1 mutant protein has increased activity as a transcription factor (Burkholder, Kurtser, and Grossman, 2001). We found that the *dnaA1* mutation also caused increased replication initiation at permissive growth temperatures. We measured replication initiation in the *dnaA1* mutant by marker frequency analysis of the origin (*ori*) and terminus (*ter*) regions of chromosomes for cells grown in defined minimal medium with glucose as a carbon source. Increased *ori/ter* typically indicates increased initiation (or decreased elongation, see below). The *dnaA1* mutant had a ∼30% increase in *ori/ter* (Table 1), consistent with an increase in initiation of DNA replication. The increase in replication initiation is likely due to an increase in the activity of the DnaA1 mutant protein at the permissive temperature, similar to its increased activity as a transcription factor (Burkholder, Kurtser, and Grossman, 2001).

**Table 1:** *ori/ter* relative to wild type

| Strain | Relevant genotype | *ori/ter*<sup>1</sup> |
| --- | --- | --- |
| AG1866 | Wild type | 1.00 ± 0.04 |
| KPL2 | *dnaA1* | 1.28 ± 0.12 |
| MEA64 | $\Delta yabA$ | 1.38 ± 0.05 |
| CAL2320 | *dnaA1* $\Delta yabA$ | 1.84 ± 0.26 |
| MEA537 | *dnaA1* $\Delta yabA$ $\Delta nrdR$ | 0.84 ± 0.09 |
| MEA102 | *dnaA1* $\Delta yabA$ *relA102*($\Delta 264-295$)<sup>2</sup> | 1.04 ± 0.31 |
| MEA370 | *dnaA1* $\Delta yabA$ *PdnaC*(T7) | 1.11 ± 0.02 |
| MEA303 | *dnaA1* $\Delta yabA$ $\Delta cshA$ | 1.55 ± 0.21 |
| MEA323 | *dnaA1* $\Delta yabA$ $\Delta ccrZ$ | 1.36 ± 0.28 |

<sup>1</sup> Strains were grown at 37°C in minimal glucose medium, and exponentially growing cells were collected to isolate genomic DNA. qPCR was used to determine marker frequency of the origin/terminus. *ori/ter* ratios are normalized to wild type (wt=1). A minimum of 3 biological replicates were included for average and standard error of the mean.
<sup>2</sup> MEA102 is the originally isolated suppressor mutation in *relA*. All other strains are reconstructed in a clean mutant background.

Null mutations in *yabA* also cause an increase in replication initiation (Hayashi *et al*., 2005; Goranov *et al*., 2009). One possible explanation is that *dnaA1* prevents or reduces the ability of YabA to inhibit replication initiation. For example, it seemed possible that the DnaA1 mutant protein might not interact normally with YabA. If true, then a *dnaA1 yabA* double mutant would have the same phenotype as a *yabA* single mutant.

We found that a *dnaA1 ΔyabA* double mutant had a much more severe phenotype than either single mutant. We introduced a *yabA* null mutation (*ΔyabA*::*spc*, simply referred to as *ΔyabA*) into a *dnaA1* mutant. Consistent with previous reports (Hayashi *et al*., 2005; Goranov *et al*., 2009), we found that *yabA* mutant cells had an approximately 40% increase in *ori/ter* when grown in defined minimal glucose medium (Table 1). We found that the *ΔyabA dnaA1* double mutant had a significant growth defect in defined minimal medium. It had a doubling time of 98 minutes compared to 50 minutes for the isogenic wild type strain. In addition, the *ΔyabA dnaA1* double mutant made small colonies on agar plates with minimal medium. In defined minimal medium, the *ΔyabA dnaA1* mutant had an *ori/ter* ratio that was ∼80-85% greater than that of wild type cells (Table 1), indicating that the effects of *dnaA1* and Δ*yabA* are roughly additive. Based on these results, we conclude that the primary defect of the *dnaA1* mutant is not loss of interaction between DnaA1 and YabA.

Too much replication initiation can lead to collapse of replication forks (Magdalou *et al*., 2014) and induction of the *recA-*dependent SOS response. We found that the SOS response was induced in the *ΔyabA dnaA1* double mutant growing in defined minimal glucose medium. We used RT-qPCR to measure mRNA levels of the DNA damage-inducible gene *dinC* (Gillespie and Yasbin, 1987; Goranov *et al*., 2006; Love, Lyle, and Yasbin, 1985; Cheo, Bayles, and Yasbin, 1991) relative to that of so-called house-keeping genes *rpoD* (*sigA*) and *gyrA*. During growth in minimal medium, levels of *dinC* mRNA were increased approximately 4-fold in the *ΔyabA dnaA1* double mutant, relative to that in wild type cells (Fig. 1). These results indicate that there is indeed DNA damage in the Δ*yabA dnaA1* double mutant. We infer that this is likely due to replication fork collapse caused by too much replication initiation. This could explain the decreased apparent growth rate of cells in liquid medium.

**Figure 1.**
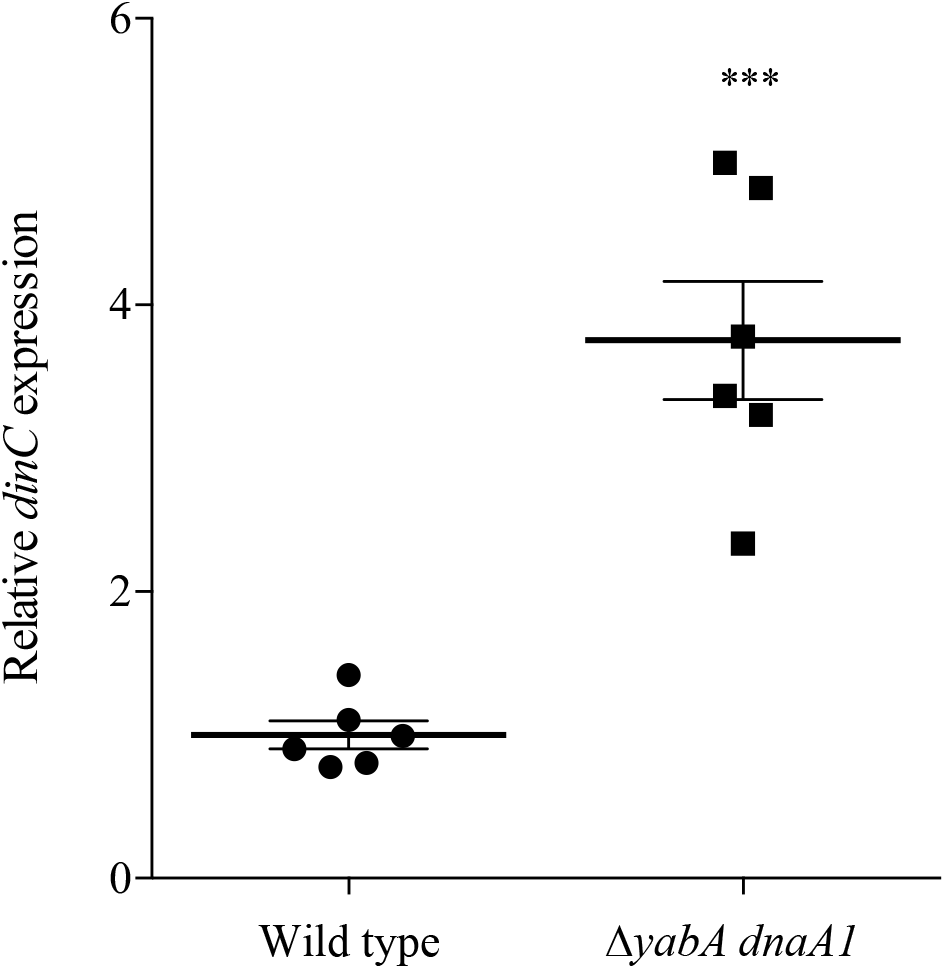
SOS-response is elevated in *ΔyabA dnaA1* compare to wild type. Strains were grown at 37°C in minimal glucose medium to mid-exponential phase and collected to isolate RNA. cDNA was synthesized using reverse transcriptase and RT-qPCR was used to measure expression of *dinC*, a gene activated during the DNA damage response. *dinC* expression was normalized to housekeeping genes, *sigA* and *gyrA*. Error bars represent standard error of the mean of 6 biological replicates. Wild type (AG1866); Δ*yabA dnaA1* (CAL2320).

In addition to the compromised growth in minimal medium, we found that the *ΔyabA dnaA1* double mutant did not grow in LB liquid medium nor did it form colonies on LB agar plates. Thus, the Δ*yabA dnaA1* double mutant had a conditional lethal phenotype that was dependent on the growth medium. Based on the phenotypes in minimal medium, the known effects of over-replication, and the increase in the frequency of replication initiation in rich compared to minimal medium, we suspect that the lethal phenotype in LB medium is due to over-initiation of replication. This conditional phenotype (able to grow in minimal but not rich medium) provided an opportunity to select for suppressor mutations that would enable survival on rich medium. We anticipated that at least some of these suppressors would affect replication initiation or replication elongation.

### Isolation of suppressors of the synthetic lethal phenotype caused of the Δ*yabA dnaA1* double mutant

We isolated 45 independent suppressors of Δ*yabA dnaA1* double mutant that restored the ability to grow on LB agar plates. Briefly, 45 independent cultures of the *ΔyabA dnaA1* double mutant (strain CAL2320) were grown in defined minimal medium and then an aliquot from each was plated on LB agar and grown at 37°C. Suppressor mutants (revertants) that were able to grow arose at a frequency of approximately 1 per 10^5^ cells. To ensure independent suppressor mutations, a single colony was chosen from each plate for further analyses. We were most interested in pseudo-revertants that were not in *dnaA*. Since we used a deletion-insertion of *yabA*, it was not possible to get suppressor mutations in *yabA*. To eliminate suppressors with mutations in *dnaA*, we amplified by PCR and sequenced *dnaA* from each of the suppressor strains. Five of the suppressors had at least one additional mutation in *dnaA* and these were not analyzed further.

We chose 30 independent suppressors for whole genome sequencing to determine the gene altered in each suppressor. Most strains had multiple mutations, but comparing them enabled us to focus on a handful of extragenic suppressors. There were several classes of suppressors that each had a mutation in the same gene (Table 2). For example, we isolated 10 independent mutants that had the same mutation in the promoter region of *dnaC*, (encoding the replicative helicase). Two or more independent mutations were also found in each of *nrdR, relA, cshA*, and *ytmP* (Table 2). These results indicated that mutations in these genes were likely responsible for enabling the *ΔyabA dnaA1* double mutant to grow on LB agar plates.

**Table 2:** Mutations that suppress the conditional synthetic lethal phenotype of Δ*yabA dnaA1*

| Gene | Occurrences | Nature of mutation |
| --- | --- | --- |
| *nrdR* | 1 | $\Delta 329$ bp; deletion of 329 bp |
|  | 1 | change of arg to leu at amino acid 12 (R12L) |
| *relA* | 1 | Nonsense mutation at amino acid S600 |
| | 1 | $\Delta 264$-295 AA in the synthetase domain of *relA*<br>(734 total AA) |
| *dnaC* | 10 | Mutation in the promoter region of *dnaC* |
| *cshA* | 1 | T27I; first recA-like domain |
|  | 2 | H312Y; second recA-like domain |
| | 1 | $\Delta 169$ bp in c-terminal domain |
| *ytmP* | 1 | R61P |
|  | 1 | A17V |

To confirm that these mutations (Table 2) were responsible for suppression of the *ΔyabA dnaA1* double mutant, we constructed defined mutations in each gene and tested each in combination with the Δ*yabA* and *dnaA1* mutations (Table 3). As the Δ*yabA dnaA1* mutant is substantially less competent and can accumulate suppressor mutations, these defined alleles were first moved into a *dnaA1* single mutant. The *yabA* deletion was moved in last, and transformants (*ΔyabA*::*spc*) were selected for resistance to spectinomycin on agar plates with defined minimal medium to minimize selection of more suppressor mutations. These reconstructed mutants were then tested for growth in rich medium (LB) to confirm the mutations were sufficient for suppression. Each of the different classes of suppressors had a decreased *ori/ter* ratio in minimal medium compared to the Δ*yabA dnaA1* double mutant parent (Table 1), consistent with the notion that the conditional lethal phenotype of the parent was due to over-initiation of replication.

**Table 3:** *B. subtilis* strains used.

| Strain | Genotype (reference) |
| --- | --- |
| AG174 | *trpC2 pheA1* (Perego *et al.*, 1988; Smith, Goldberg, and Grossman, 2014) |
| AG1866 | *trpC2 pheA1* Tn917ΩHU163 (*mls*)<sup>1</sup> |
| KPL2 | *trpC2 pheA1 dnaA1*-Tn917ΩHU163 (*mls*) (Burkholder *et al.*, 2001) |
| CAL2320 | *trpC2 pheA1 ΔyabA::spc dnaA1*-Tn917ΩHU163 (*mls*) |
| MEA73<sup>2</sup> | *trpC2 pheA1 ΔyabA::spc dnaA1*-Tn917ΩHU163 (*mls*) *PdnaC(T7)* |
| MEA64 | *trpC2 pheA1 ΔyabA::spc* Tn917ΩHU163 (*mls*) |
| MEA102<sup>2</sup> | *trpC2 pheA1 ΔyabA::spc dnaA1*-Tn917ΩHU163 (*mls*) *relA102(Δ264-295)* |
| MEA187 | *trpC2 pheA1 ΔnrdR::kan* |
| MEA250 | *trpC2 pheA1 ΔcshA::kan* |
| MEA303 | *trpC2 pheA1 ΔyabA::spc dnaA1*-Tn917ΩHU163 (*mls*) *ΔcshA::kan* |
| MEA304 | *trpC2 pheA1 ΔytmP::kan* |
| MEA323 | *trpC2 pheA1 ΔyabA::spc dnaA1*-Tn917ΩHU163 (*mls*) *ΔytmP::kan* |
| MEA359 | *trpC2 pheA1 ΔpurA::kan* |
| MEA360 | *trpC2 pheA1 ycgO::Pspank-dnaC-cat* |
| MEA361 | *trpC2 pheA1 ycgO::Pspank-dnaC-cat ΔdnaC::kan* |
| MEA362 | *trpC2 pheA1 PdnaC(T7)* |
| MEA370 | *trpC2 pheA1 ΔyabA::spc dnaA1*-Tn917ΩHU163 (*mls*) *PdnaC(T7)* |
| MEA415 | *trpC2 pheA1 polC::Pspank-polC-spc* |
| MEA537 | *trpC2 pheA1 ΔyabA::spc dnaA1*-Tn917ΩHU163 (*mls*) *ΔnrdR::kan* |

<sup>1</sup> The Tn917ΩHU163 is in *dck* and linked to *dnaA*.<sup>2</sup> These were the original strains isolated from the suppressor screen. All other strains were constructed in a clean background.

### Characterization of suppressor mutants

We characterized suppressors with mutations in *relA, nrdR, dnaC, cshA*, and *ytmP*, with extensive analysis of the mutations affecting *dnaC* (encoding the replicative DNA helicase). In all cases, we measured the relative copy number of the *oriC* region to that of *terC* (*ori* to *ter* ratios or *ori/ter*) in the suppressor strains compared to that of the Δ*yabA dnaA1* double mutant (Table 1; described below). Although *ori/ter* is frequently used as a readout of DNA replication initiation, it is important to note that changes in *ori/ter* could be due to changes in replication initiation or changes in replication elongation. An increase in the overall rate of replication elongation would result in an increase in the copy number of the terminus region, thereby causing a reduction in the marker frequency ratio of *ori*/*ter*. Below, we describe results that indicate that some suppressor mutations affect replication initiation and others affect replication elongation.

### Mutations in *relA* and *nrdR* stimulate replication elongation

#### relA

We isolated several mutations in *relA*. The *relA* gene product interacts with the ribosome and is responsible for most of the synthesis and hydrolysis of (p)ppGpp in *B. subtilis* (Wendrich and Marahiel, 1997; Srivatsan and Wang, 2008; Nanamiya *et al*., 2008; Kriel *et al*., 2014). *B. subtilis*, RelA has three domains, one for synthesis and one for hydrolysis of (p)ppGpp, and a third domain required for interaction with the ribosome (Wendrich and Marahiel, 1997). The suppressor mutations affected either the (p)ppGpp synthetase or ribosome binding domain, and none were in the hydrolase domain (Table 2). One of these mutants (MEA102) had an in-frame deletion that removes the conserved synthetase active site residue Asp^264^ (Kriel *et al*., 2014), indicating that this mutation causes a decrease in (p)ppGpp synthesis. Decreased levels of (p)ppGpp likely stimulate replication elongation in *B. subtilis* through reduced inhibition of the primase, DnaG (Wang, Sanders, and Grossman, 2007). The presumed decrease in (p)ppGpp in the *relA* suppressor mutants would therefore indicate an increase in the processivity of the replication fork, which could cause the observed decrease in *ori/ter*.

We characterized one of the originally isolated suppressor mutants (MEA102) that contains a mutation in *relA* (*relA102*), that should inactivate the synthetase domain (Table 1; MEA102). We used the original mutant because we were unable to reconstruct an appropriate synthetase null in the Δ*yabA dnaA1* background, largely due to lack of competence of the *relA* mutant and the Δ*yabA dnaA1* mutant. The *relA102* Δ*yabA dnaA1* suppressor mutant had a decrease in *ori/ter* in cells growing in defined minimal medium compared to the parent *ΔyabA dnaA1* double mutant (Table 1). This could indicate a decrease in replication initiation or an increase in replication elongation (or both).

We measured the rate of DNA synthesis by pulse-labeling cells with ^3^H-thymidine and measuring incorporation into DNA (Methods). We found that there was an approximately 50% increase in the rate of DNA synthesis in the suppressor mutant (*relA102 ΔyabA dnaA1*) compared to that in the parent (*ΔyabA dnaA1*) (Fig. 2A) in cells that were in mid-exponential growth in defined minimal medium. The increased rate of DNA synthesis in the suppressor mutant (Fig. 2A), combined with the decrease in *ori*/*ter* (Table 1) indicated that the *relA* mutation primarily caused an increase in replication elongation (and not a decrease in replication initiation). This effect is most likely because of an increase in the activity of DNA primase due to decreased inhibition by (p)ppGpp.

**Figure 2.**
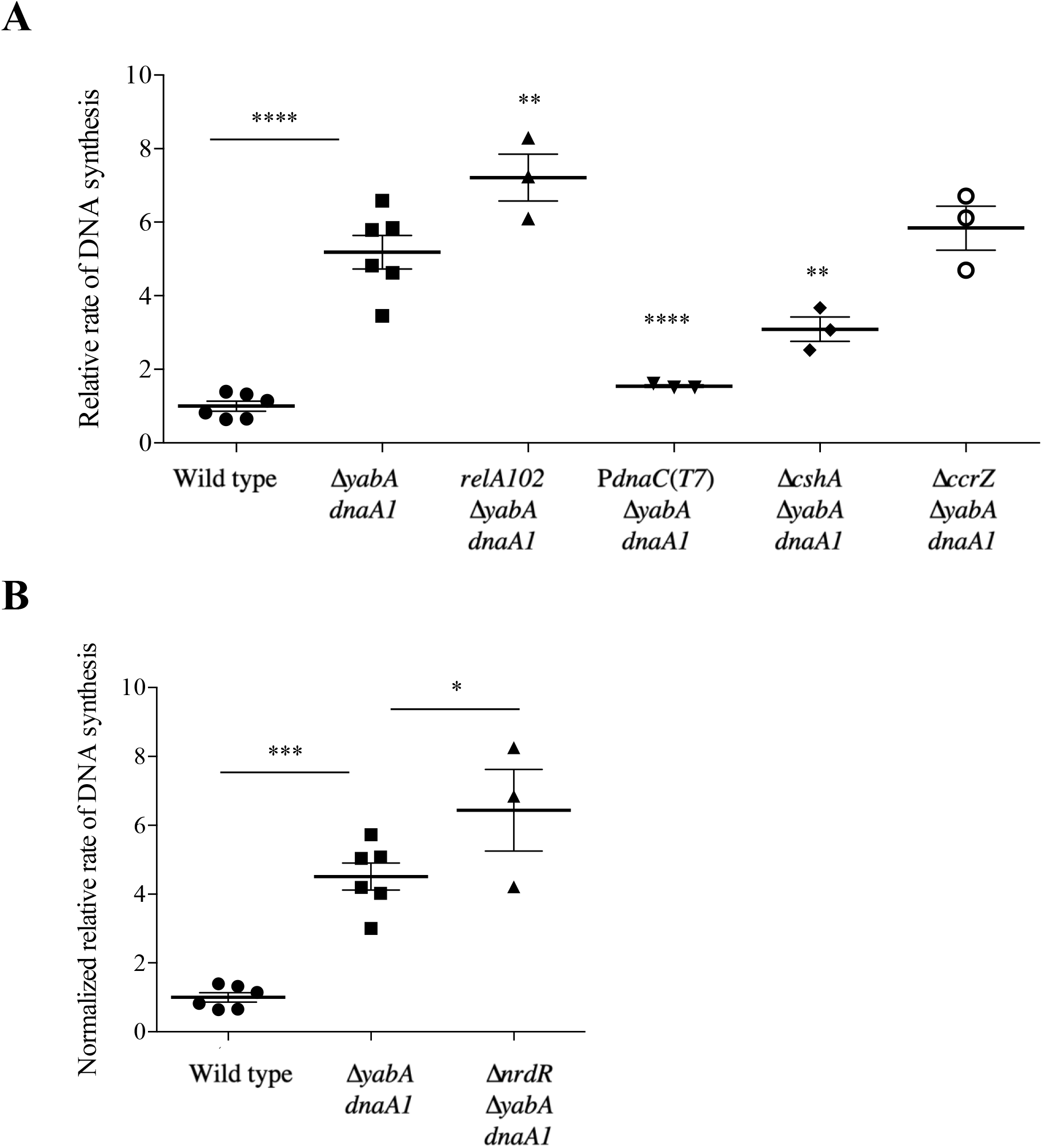
Relative rate of DNA synthesis. Strains were grown at 37°C in minimal glucose medium to mid-exponential phase. Relative rate of DNA synthesis was measured (incorporation of ^3^H-thymidine into DNA relative to wild type). **A**. Wild type (AG1866); Δ*yabA dnaA1* (CAL2320); Δ*yabA dnaA1 relA102* (MEA102); Δ*yabA dnaA1* P*dnaC*(*T7*) (MEA370); Δ*yabA dnaA1* Δ*cshA* (MEA303); Δ*yabA dnaA1* Δ*ccrZ* (MEA323). MEA102 is the originally isolated suppressor mutation in *relA*. All other strains are deletions reconstructed in a clean mutant background. Error bars represent standard error of the mean for at least 3 biological replicates. Significant differences compared to the parent strain (CAL2320) are indicated (P < 0.05). **B**. Wild type (AG1866); Δ*yabA dnaA1* (CAL2320); Δ*yabA dnaA1* Δ*nrdR* (MEA537). ^3^H-thymidine incorporation was normalized to the mean (of at least 3 biological replicates) cellular levels of dTMP in each strain as determined by mass spectrometry (Table 4; Methods). Error bars represent standard error of the mean of the normalized ^3^H-thymidine incorporation for at least 3 biological replicates. Significant differences compared to the parent strain (CAL2320) are indicated (P < 0.05).

#### nrdR

Two independent suppressor mutations were isolated in *nrdR* (negative regulator of ribonucleotide reductase genes), one of which was almost a complete deletion of the gene (Table 2), indicating that loss of *nrdR* likely suppressed the Δ*yabA dnaA1* double mutant. To confirm that *nrdR* was responsible for suppression, we constructed a deletion of *nrdR* and made the triple mutant Δ*nrdR* Δ*yabA dnaA1*. This mutant was indeed able to grow on LB agar, confirming that Δ*nrdR* is sufficient for suppression of the lethality of the *ΔyabA dnaA1* mutant.

**Table 4:** Δ*nrdR* has increased pools of dTMP.

| Strains<sup>1</sup> | Relative dTMP<sup>2</sup> |
| --- | --- |
| Wild type | 1.00 ± 0.03 |
| $\Delta yabA dnaA1$ | 0.87 ± 0.17 |
| $\Delta nrdR \Delta yabA dnaA1$ | 3.73 ± 0.96 |

<sup>1</sup> Wild type (AG1866); $\Delta yabA dnaA1$ (CAL2320); $\Delta nrdR \Delta yabA dnaA1$ (MEA537).
<sup>2</sup> Samples grown at 37°C in minimal glucose medium to mid-exponential phase were collected for metabolomic analysis using 40% methanol, 40% acetonitrile, 20% water with 0.1M formic acid, and 500 nM each of 17 isotopically-labeled amino acids for extraction buffer (see Methods). Values represent the mean and standard error of the mean of at least three biological replicates normalized to wild type.

We measured the *ori/ter* ratio in the Δ*nrdR* Δ*yabA dnaA1* strain and observed a significant decrease compared to that of the Δ*yabA dnaA1* double mutant (Table 1). Although *ori/ter* is frequently used as a readout of DNA replication initiation, we again note that a decrease in *ori/ter* could be due to either a decrease in replication initiation or an increase in replication elongation (or both). Because of the known function of NrdR, we anticipated that the suppressor had an increase in replication elongation.

NrdR controls levels of the nucleotide pools and a deletion of *nrdR* is known to cause an increase in the overall levels of nucleotides in the cell (Grinberg *et al*., 2006; Torrents *et al*., 2007). Presumably, this effect could help stimulate replication elongation in the Δ*yabA dnaA1* strain, which over-initiates and therefore might require higher levels of nucleotide pools for processive elongation.

Whereas incorporation of ^3^H-thymidine in pulse-labeling experiments can be used to measure the rate of DNA replication, incorporation is also influenced by the pool size of thymidine in cells. In many analyses and comparisons, there are no expectations that pool sizes will vary between strains. However, since loss of *nrdR* is known to affect nucleotide biosynthesis (Grinberg *et al*., 2006; Torrents *et al*., 2007), we measured pool sizes to correct for possible changes in specific activity of ^3^H-thymidine inside cells.

As expected, the concentration of dTMP in the *nrdR* mutant strains was increased. We used mass spectrometry to measure levels of dTMP in an *nrdR* null mutant compared to wild type or the parent *ΔyabA dnaA1* strains (Table 4). Cells were grown in defined minimal medium at 37°C and samples were taken for analysis (Methods). There was an increase in the concentration of dTMP of approximately 3-4-fold in the *ΔnrdR* mutants. Using this information, we could now compare rates of DNA synthesis between cells with different intracellular concentrations of thymidine nucleotides.

We found that there was an increase in the rate of DNA synthesis in the Δ*nrdR ΔyabA dnaA1* triple mutant compared to the *ΔyabA dnaA1* (*nrdR*^+^) parent. Cells were grown to exponential phase in defined minimal medium at 37°C and pulse labeled with ^3^H-thimidine (Methods, and as above). Taking into account the differences in the intracellular concentration of thymidine nucleotides (Table 4), there was an approximately 40% increase in the rate of DNA replication in the *ΔnrdR ΔyabA dnaA1* triple mutant compared to the *nrdR*^+^ double mutant (*ΔyabA dnaA1*) parent (Fig. 2B). Together with the decrease in *ori/ter*, these data indicate that the Δ*nrdR* mutation suppresses Δ*yabA dnaA1* primarily by stimulating replication elongation and not by decreasing replication initiation.

### Suppressor mutations in the promoter region of *dnaC*, the gene encoding the replicative helicase

Ten of the suppressors had a single base pair deletion in the promoter region of *dnaC*, the gene that encodes the replicative DNA helicase. The mutation P*dnaC*(*T7*) changes a run of eight thymine’s to seven, resulting in a decrease in the spacing between the -10 and -35 regions of the promoter driving expression of *dnaC*. This decrease in spacing likely causes a decrease in promoter activity.

To confirm that this mutation was responsible for suppression of the *ΔyabA dnaA1* mutant phenotype, we constructed a single base pair deletion {P*dnaC*(*T7*)} and made the triple mutant P*dnaC*(*T7*) Δ*yabA dnaA1*. This reconstructed mutant {P*dnaC*(*T7*) Δ*yabA dnaA1*} grew in rich (LB) medium, confirming that the single base pair deletion in the promoter region of *dnaC* was sufficient to suppress the conditional lethal phenotype of the *ΔyabA dnaA1* double mutant. Below we describe experiments with the P*dnaC*(*T7*) mutants to determine the mechanism of suppression.

We hypothesized that the P*dnaC*(*T7*) mutation likely caused a change in expression of *dnaC*, and consequently a change in the amount of the DnaC protein (helicase). We measured the amount of helicase protein in cells grown in minimal medium with glucose, using Western blots and probing with antibodies to DNA helicase. Both the wild type and Δ*yabA dnaA1* had roughly the same amount of the replicative helicase, whereas the Δ*yabA dnaA1* P*dnaC*(*T7*) suppressor had ∼50% of wild type levels (Fig. 3A). A similar decrease in helicase protein was observed for the single mutant, i.e. P*dnaC*(*T7*) in a wild-type (*dnaA*+ *yabA*+) background (Fig 3B). These data indicate the P*dnaC*(*T7*) mutation caused a decrease in the amount of the replicative helicase (DnaC).

**Figure 3.**
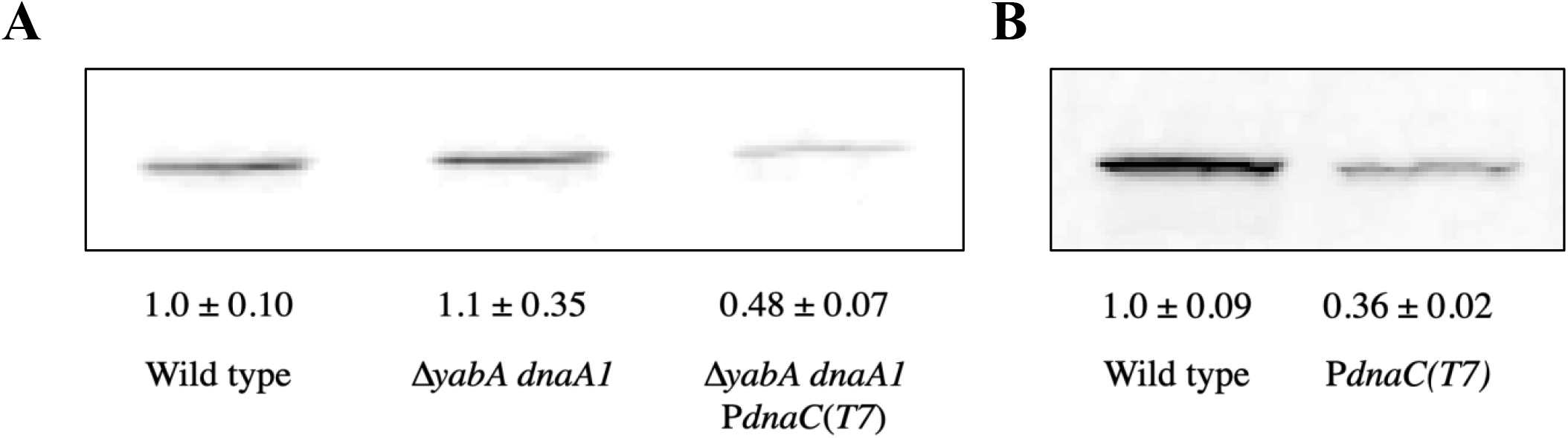
P*dnaC*(*T7*) causes decreased levels of DnaC protein. Strains were grown at 37°C in minimal glucose medium to mid-exponential phase. Levels of DnaC were determined by quantitative western blot, normalized to optical density (A_600_; Methods). One representative blot is shown and values represent the average and standard error of the mean of at least 3 biological replicates. **A**. Wild type (AG1866); Δ*yabA dnaA1* (CAL2320); Δ*yabA dnaA1* P*dnaC*(*T7*) (MEA370). **B**. Wild type (AG174); P*dnaC*(*T7*) (MEA362).

### Decreased levels of the replicative helicase are sufficient to lower DNA replication initiation under fast growth conditions

Because the replicative helicase is needed for replication initiation (and elongation), we hypothesized that the decreased levels of helicase in the P*dnaC*(*T7*) mutants might reduce replication initiation and thereby be responsible for suppressing the overinitiation observed in the Δ*yabA dnaA1* double mutant. We measured *ori/ter* ratios in the P*dnaC*(*T7*) suppressor and found that the P*dnaC*(*T7*) mutation caused a significant decrease in *ori/ter* close to the levels of wild type (minimal glucose; Table 1). To determine whether the decrease in *ori/ter* was due to decreased initiation or increased elongation, the relative rate of DNA synthesis was measured. Similar to *ori/ter* results, the P*dnaC*(*T7*) mutation caused a decrease in total DNA synthesis compared to the Δ*yabA dnaA1* parent (minimal glucose; Fig. 2A). Taken together, these data indicate that the P*dnaC*(*T7*) mutation causes a decrease in DNA replication initiation in a Δ*yabA dnaA1* background, presumably due to the decrease in the amount of the replicative helicase.

We found that the P*dnaC*(*T7*) mutation also caused a decrease in *ori/ter* in otherwise wild type cells under conditions of rapid growth in rich medium (LB). During rapid growth in LB medium, the *ori/ter* of the P*dnaC*(*T7*) mutant had a ∼20% decrease compared to the isogenic wild type strain (Fig. 4A). During slower growth in minimal glucose medium, the P*dnaC*(*T7*) mutant had no detectable change in *ori/ter* (Fig. 5), despite the fact that it contained decreased amounts of the replicative helicase (Fig. 3B). These results indicate that the decrease in the amount of the replicative helicase only has an effect on initiation when the cells are experiencing high rates of DNA replication initiation, such as fast growth (LB) or in the Δ*yabA dnaA1* mutant.

**Figure 4.**
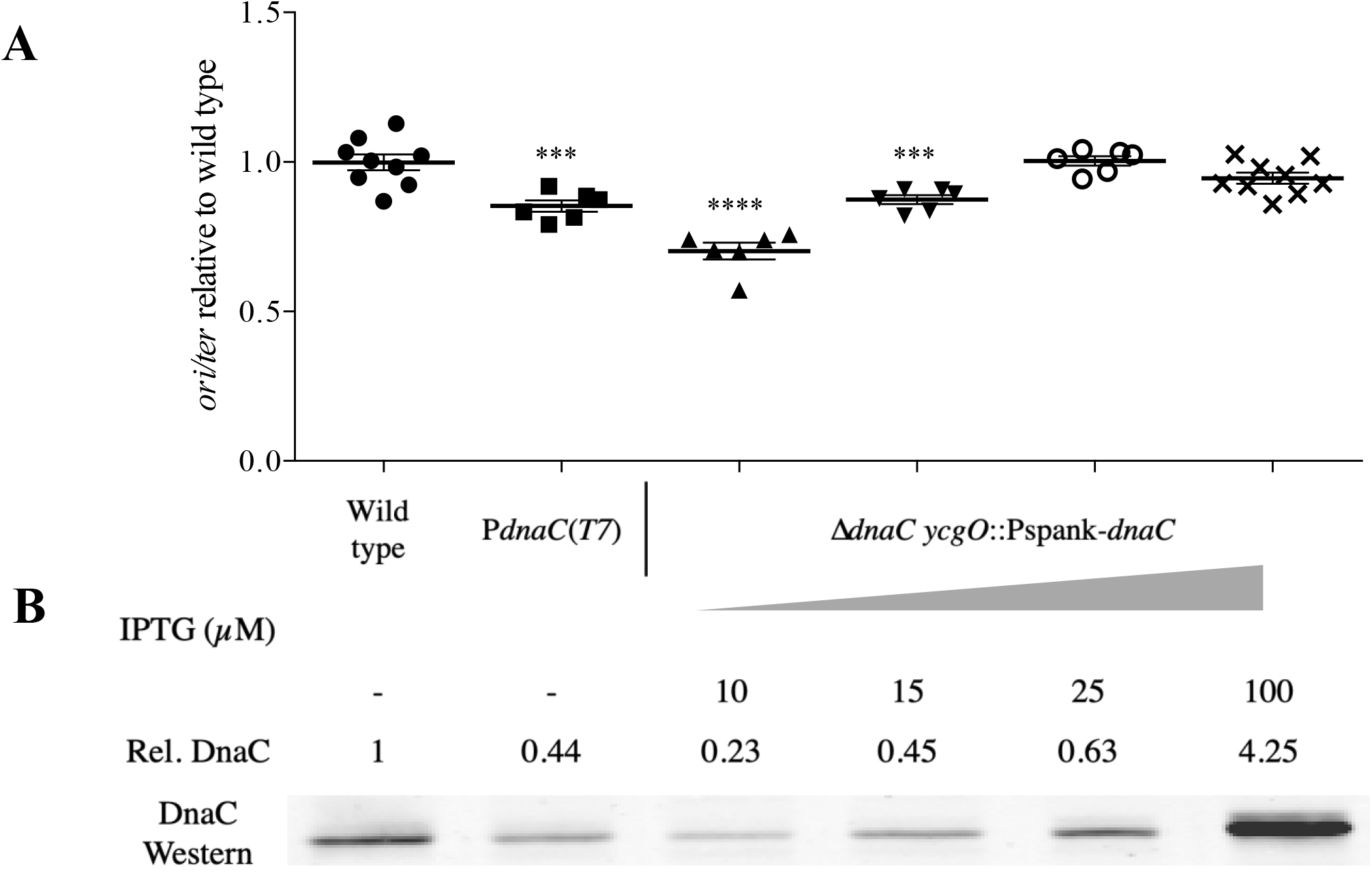
Decreased levels of DnaC are sufficient to decrease *ori/ter*. Strains were grown at 37°C in LB to mid-exponential phase. Expression from Pspank-*dnaC* was induced by addition of varied levels of IPTG (10 µM-100µM). Wild type (AG174); P*dnaC*(*T7*) (MEA362); Δ*dnaC ycgO*::Pspank-*dnaC* (MEA361). **A**. Samples were taken for genomic DNA isolation and used for qPCR to measure *ori/ter* as in Table 1. Error bars represent standard error mean for at least 6 biological replicates. Significant differences compared to the wild type (AG174) are indicated (P < 0.05). **B**. At the same time, samples were taken to measure DnaC protein with quantitative western blots as in Figure 3. This is a representative experiment but was repeated with 3 biological replicates.

**Figure 5.**
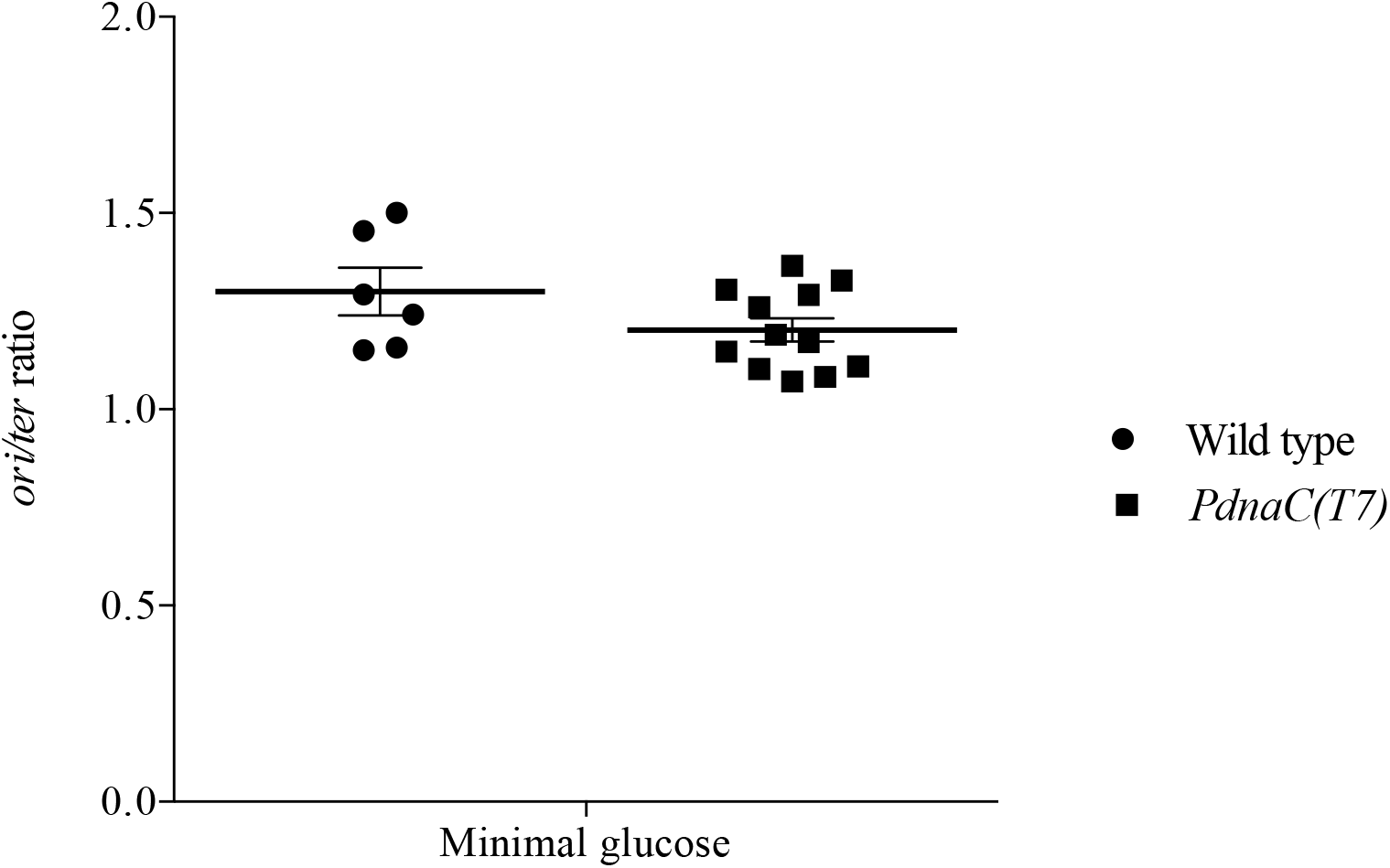
P*dnaC*(*T7*) does not affect *ori/ter* in slow growth (minimal glucose) conditions. Strains were grown at 37°C in minimal glucose medium to mid-exponential phase and cells were collected to isolate genomic DNA. qPCR was used to measure *ori/ter* as in Table 1. Error bars represent standard error of the mean for at least 6 biological replicates. Wild type (AG174); P*dnaC*(*T7*) (MEA362).

If the decreased initiation phenotype of the P*dnaC*(*T7*) mutation is really due to altered levels of helicase in the mutant, then we should be able to reproduce the phenotype by varying expression of *dnaC* with a controllable promoter. To vary the amount of the replicative helicase in cells, we fused *dnaC* to the LacI-repressible-IPTG-inducible promoter Pspank (Pspank-*dnaC*). Cells required IPTG for growth, and by growing in varied concentrations of IPTG different amounts of replicative helicase were obtained, ranging from ∼25 to 425% of levels in wild type cells (Fig. 4B). We found that there was a decrease in *ori/ter* as the levels of DnaC decreased. There was a significant decrease in *ori/ter* when levels of DnaC were decreased below roughly 50% that of wild type (Fig. 4A). At higher levels of DnaC the *ori/ter* ratio was essentially unchanged, even when DnaC was overexpressed ∼4-fold over wild type levels.

### Decreasing levels of PolC does not decrease DNA replication initiation

In order to show that the decrease in DNA replication initiation was specific to decreased levels of the helicase, and not a general trend among other proteins involved in DNA replication initiation, we varied expression levels of DNA polymerase, *polC*, and measured *ori/ter*. We constructed an IPTG-inducible version of *polC* and measured *ori/ter* under a range of induction levels. Unlike the titration of *dnaC*, we did not observe a decrease in *ori/ter* as the levels of *polC* decreased (Fig. 6). In fact, at the lowest level of induction we observed an increase in *ori/ter*. This agrees with a previous report that observed a decrease in elongation upon depleting levels of *polC* but no effect on initiation (Dervyn *et al*., 2001). At all other levels of induction of *polC ori/ter* was indistinguishable from that of wild type (Fig. 6).

**Figure 6.**
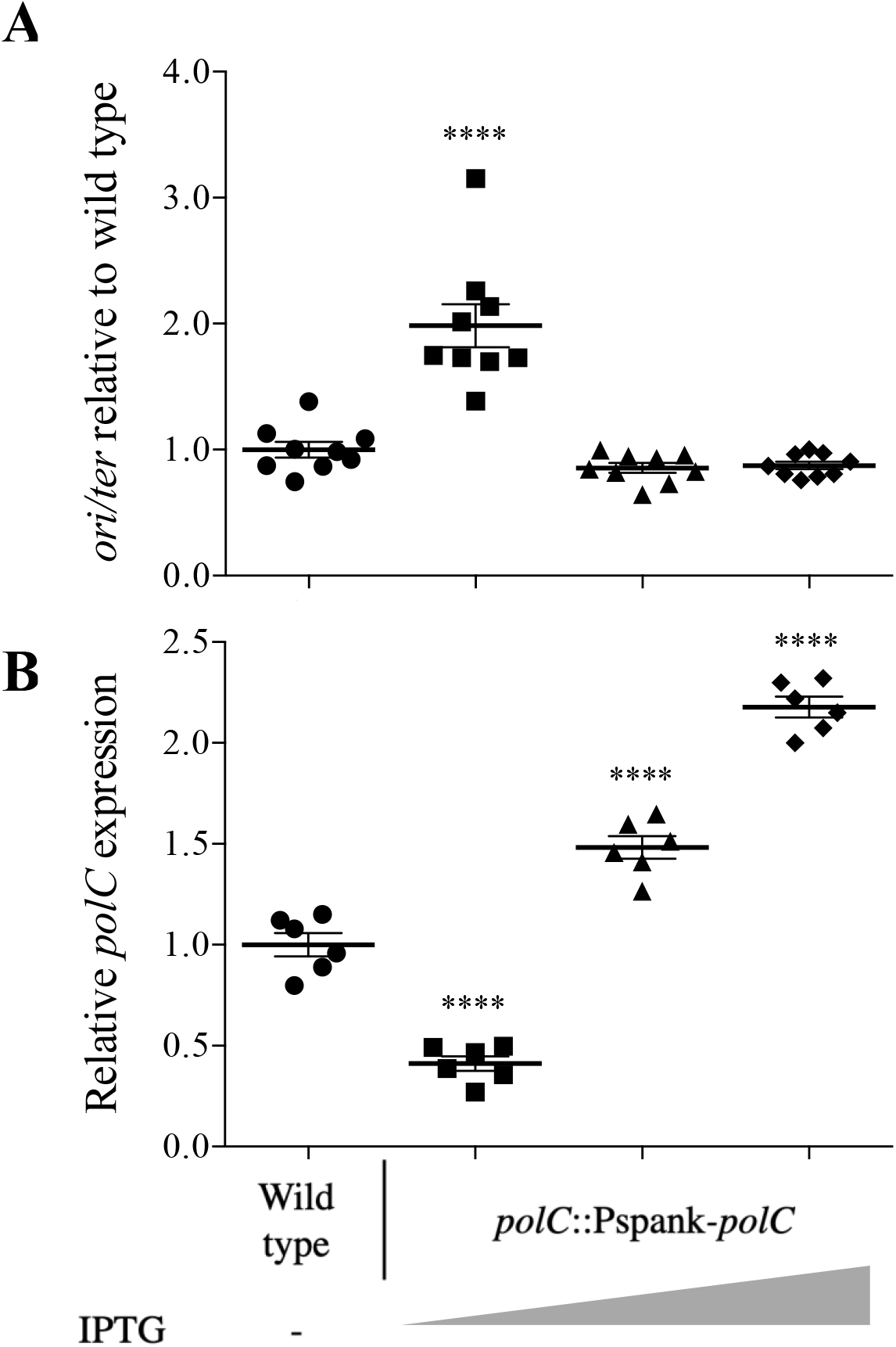
Decreased levels of *polC* does not decrease *ori/ter*. Strains were grown at 37°C in minimal glucose medium to mid-exponential phase and cells were collected to isolate RNA and genomic DNA. Expression from Pspank-*polC* was induced by addition of varied levels of IPTG. Error bars reflect standard error mean of at least 6 biological replicates. Wild type (AG174); *polC*::Pspank-*polC* (MEA415). Significant differences compared to the wild type (AG174) are indicated (P < 0.05). **A**. gDNA was used for qPCR to measure *ori/ter* as in Table 1. Error bars represent standard error of the mean of 9 biological replicates. **B**.RT-qPCR was used to measure expression of *polC* as in Figure 1. *polC* expression was normalized to housekeeping genes, *sigA* and *gyrA*. Error bars represent standard error of the mean of 6 biological replicates.

Although the suppressor screen reported here was not saturated, there were no mutations isolated in other genes involved in DNA replication (apart from *dnaA*). These data indicate that the decrease in replication initiation may be specific to the decrease in levels of the helicase, and not a general phenomenon that applies to other proteins involved in DNA replication initiation.

### Suppressors in *cshA* and *ytmP*

#### cshA

Several mutations were isolated in *cshA*, a cold-shock RNA helicase associated with the RNA degradosome, and a deletion of *cshA* was sufficient to suppress the growth phenotype in the Δ*yabA dnaA1* background. Similar to the other suppressors isolated, Δ*cshA* Δ*yabA dnaA1* had a decrease in both *ori/ter* and the rate of DNA synthesis compared to Δ*yabA dnaA1*, indicating that there was likely a decrease in DNA replication initiation (Table 1). Nucleotide pools in the *cshA* mutants were not elevated compared to those in wild type cells (92% ± 15%) or the *yabA dnaA1* mutant (58% ± 5%). Based on these results, we conclude that suppression by the *cshA* mutations was largely due to a decrease in replication initiation.

Given the role of *cshA* in mRNA turnover and ribosome biogenesis, we hypothesized that there may be a change in stability of certain mRNAs or the amount of one or more proteins associated with DNA replication. Given that we determined a decrease in expression of *dnaC*, encoding the replicative helicase, was sufficient to decrease DNA replication initiation (above), we measured levels of *dnaC* mRNA in a Δ*cshA* mutant using RT-qPCR. We found that there was an approximately 50% decrease in the amount of *dnaC* mRNA in Δ*cshA* (Fig. 7B). This may contribute to the decrease in *ori/ter* but is likely not the sole factor because there is a greater decrease in *ori/ter* in Δ*cshA* than in the P*dnaC*(*T7*) mutant in otherwise wild type cells grown in rich medium (Fig. 7A, Fig. 4A). Additionally, Δ*cshA* has a much slower growth rate compared to wild type, with Δ*cshA* having a doubling time of 51 minutes compared to 21 minutes for wild type in LB medium. This could contribute to the decrease in *ori/ter* but may also decrease the amount of DnaC needed for DNA replication initiation. It seems likely that suppression of the *dnaA1 yabA* mutant is due to more than one effect of loss of *cshA*.

**Figure 7.**
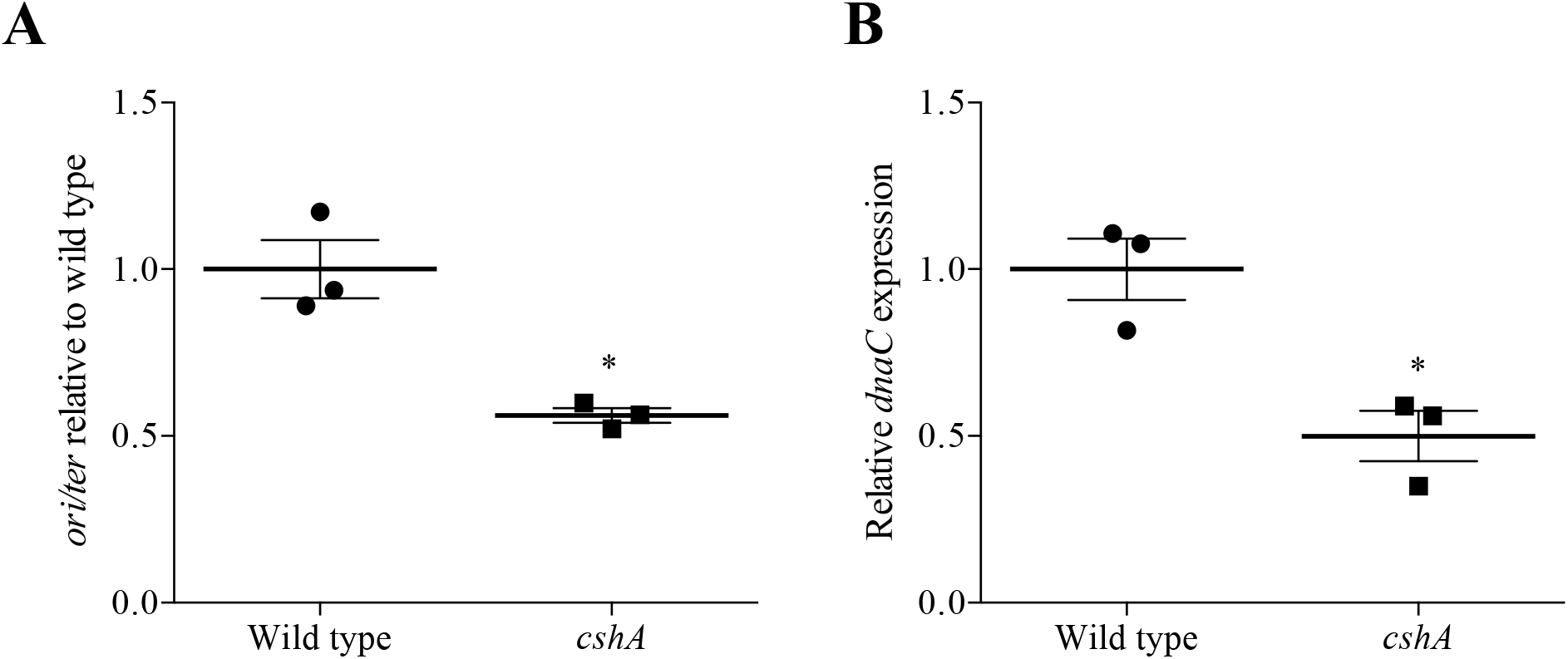
*dnaC* expression and *ori/ter* are decreased in Δ*cshA*. Strains were grown at 37°C in LB to mid-exponential phase and cells were collected to isolate RNA and genomic DNA. Error bars reflect standard error of the mean of 3 biological replicates. Wild type (AG174); Δ*cshA* (MEA250). Significant differences compared to the wild type (AG174) are indicated (P < 0.05). **A**.gDNA was used for qPCR to measure *ori/ter* as in Table 1. **B**.RT-qPCR was used to measure expression of *dnaC* as in Figure 1. *dnaC* expression was normalized to housekeeping genes, *sigA* and *gyrA*.

#### ytmP (ccrZ)

We isolated several mutations in a previously uncharacterized gene, *ytmP*, now called *ccrZ* (Gallay *et al*., 2019). The *ccrZ* gene product has regions of similarity to choline and enthanolamine kinases (Zimmermann *et al*., 2018). We constructed a null mutation in *ccrZ* and this mutation was sufficient to suppress the conditional lethal phenotype of the Δ*yabA dnaA1* mutant. The *ΔccrZ* mutation caused a decrease in *ori/ter* relative to that of the *ΔyabA dnaA1* parent (Table 1), indicating either a decrease in replication initiation and/or an increase in replication elongation. We found that DNA synthesis in the suppressor *ccrZ ΔyabA dnaA1* triple mutant had virtually no change in DNA synthesis relative to that in the parent *ΔyabA dnaA1* double mutant (Fig. 2A). Based on these results, we conclude that loss of *ccrZ* may suppress the *ΔyabA dnaA1* mutant through multiple mechanisms. It is also possible that changes replication initiation frequency could cause indirect effects on elongation. A small decrease in initiation could prevent replication fork collapse, thereby allowing for more processive replication elongation.

A homolog of *B subtilis ccrZ* was identified as an essential gene in *Streptococcus pneumoniae*. In *S. pneumoniae, ccrZ* was found to couple DNA replication with cell division. The effects of *ccrZ* in *S. pneumoniae* and *B. subtilis* are described in other work (Gallay *et al*., 2019).

## Discussion

### Suppressors of too much replication initiation

Work presented here highlights the multiple ways cells can compensate for overinitiation of DNA replication. Starting with a mutant that had a conditional lethal phenotype, presumably due to overinitiation of replication, we isolated suppressors that were capable of growing under otherwise non-permissive conditions (rich medium). Suppressors were found to cause a decrease in replication initiation and/or an increase in replication elongation. One suppressor was in the promoter region of the gene for the replicative DNA helicase and this was characterized in detail.

### Decreased levels of helicase as a novel mechanism of suppression for DNA replication overinitiation in *B. subtilis*

The majority of the suppressors we identified were in the promoter region of the replicative helicase, *dnaC*, and caused a decrease in the amount of helicase in cells. We found that in conditions in which cells have a relatively high frequency of replication initiation, either in the overinitiating double mutant (Δ*yabA dnaA1*) or fast growth conditions (LB), decreasing the amount of helicase was sufficient to reduce the frequency of replication initiation. Previous work indicated that overproduction of helicase could cause an increase in replication initiation, and this effect was dependent on *yabA* (Goranov *et al*., 2009). In contrast, the decrease in replication initiation caused be reduced amount of the helicase was independent of *yabA*, at least in the *ΔyabA dnaA1* mutant. In contrast to the effects of overproducing helicase in wild type cells (Goranov *et al*., 2009), we found that overproduction of helicase had no detectable effect on replication initiation in the *ΔyabA dnaA1* mutant.

In contrast, in *E. coli*, perturbing the ratio of helicase to helicase loader proteins inhibits replication initiation and elongation (Allen and Kornberg, 1991; Skarstad and Wold, 1995; Bruning, Myka, and McGlynn, 2016). This is believed to be the result of an imbalance between the helicase and the loader proteins, which are normally at a ∼1:1 ratio. To load the helicase ring onto the DNA, *E. coli* employs a “ring breaker” mechanism where the hexameric helicase loader breaks an already formed hexameric helicase ring and loads it around the DNA (Kornberg and Baker, 1992; reviewed in Davey and O’Donnell, 2003). An imbalance between the two proteins causes inhibitory associations between the helicase and the loader: excess loader causes continual reassociation between the helicase and loader, preventing replisome progression (Allen and Kornberg, 1991; Skarstad and Wold, 1995), whereas excess helicase reduces the probability of the helicase-loader complex formation (Bruning, Myka, and McGlynn, 2016).

*B. subtilis* and some other Gram-positive bacteria use a “ring maker” mechanism where a monomeric helicase loader loads individual monomers around the DNA to create the hexameric ring required for DNA replication (reviewed in Davey and O’Donnell, 2003; Velten *et al*., 2003). Any inhibitory interactions would only affect monomer-monomer interactions, rather than inhibiting an already assembled hexameric helicase, prevent its progression. As shown above, overproduction of DnaC had no effect on *ori/ter*, indicating that a different mechanism for decreasing replication initiation is occurring in *B. subtilis*.

We suggest that levels of DnaC are limiting under high initiation conditions, causing the decrease in replication initiation. We only observed an effect under conditions with high DNA replication initiation, such as rich medium (LB) or in the overinitiating mutant background of Δ*yabA dnaA1*. During slower growth, where lower concentrations of helicase protein are required, we observed no change in DNA replication initiation. This makes sense given that higher levels of helicase are required when more DNA replication is taking place. It is also possible that under slow growth (minimal media) conditions, where wild type cells have an *ori/ter* of only about 1.2, there is not much room for this mutation to decrease replication initiation.

### Decreased replication initiation due to lower levels of a replication protein appears to be specific to the helicase, DnaC

We did not isolate suppressor mutations in other genes known to be involved in replication initiation (apart from *dnaA)*. Most proteins involved in DNA replication are essential and therefore any loss of function mutations would not be isolated in this screen. Our screen was able to isolate mutations that affect protein levels or activity, which are much less likely to occur. The P*dnaC* mutation we isolated, which occurred in the promoter region of *dnaC*, is one example of this type of mutation. The mutation that caused the decreased levels of helicase was a single basepair deletion in a run of thymines. However, we did not isolate any similar mutations in other essential DNA replication genes. This is not surprising, as these types of mutations occur very rarely and our screen was not performed to saturation. It is possible that the specific mutation in P*dnaC* occurs at higher frequency than promoter mutations in other DNA replication genes, due to the tendency of slippage by the replication machinery at poly-N tracts (reviewed in Strauss, 1999). It is unique that P*dnaC* is so easy to mutate. Other genes involved with replication initiation, such as *dnaB-dnaI* and *dnaD*, have shorter poly-N tracts in their promoters (≤5) and not in the -10 to -35 region.

It is also possible that we did not isolate mutations affecting the levels of other DNA replication proteins because these proteins are present in excess and a small decrease in levels would not affect initiation. If this is true, helicase is unique in being synthesized to roughly the required amount for fast growth and not in excess. Decreased expression of another protein involved in DNA replication, the replicative polymerase (encoded by *polC*), did not decrease DNA replication, although this does not rule out other proteins required for replication initiation.

### Mutations that increase replication elongation can suppress growth defects caused by overinitiation

The conditional lethal phenotype of the Δ*yabA dnaA1* mutant is most likely caused by replication overinitiation, leading to replication fork collapse. We isolated several mutations that suppress the overinitiation mutant by causing a decrease in replication initiation. We also isolated suppressor mutations that caused an increase in replication elongation. Increased replication elongation could help overcome the overinitiation defect by either preventing replication fork collapse, or stimulating repair of collapsed forks.

We determined that mutations in *relA*, likely resulting in a decrease in (p)ppGpp, appear to stimulate replication elongation. (p)ppGpp is known to inhibit elongation, specifically by inhibiting primase (DnaG) (Wang, Sanders, and Grossman, 2007; Maciag *et al*., 2010). DNA primase synthesizes an RNA primer required for lagging strand DNA synthesis. Decreased inhibition of primase would help increase replication elongation. This could be by stimulating the rate of elongation in the mutant background thereby preventing replication fork collapse, by enhancing replication restart from already collapsed forks, or a combination of these two mechanisms.

Similarly, we determined that a null mutation in *nrdR* also suppressed the overinitiation mutant by stimulating replication elongation. Changes in expression of *nrd* genes, which encode nucleotide reductase needed for deoxyribonucleotide biosynthesis (reviewed in (Nordlund and Reichard 2006), in relation to perturbations in DNA replication have been well documented (Augustin, Jacobson, and Fuchs, 1994; Huang, Zhou, and Elledge, 1998; Goranov *et al*., 2005), establishing a link between DNA replication and nucleotide biosynthesis. In *E. coli*, DnaA itself directly activates the *nrdAB* operon (Augustin, Jacobson, and Fuchs, 1994), indicating a positive correlation between increased DNA replication initiation and dNTPs. In *B. subtilis*, expression of the *nrdEF* operon (regulated by *nrdR*) increases upon replication fork arrest (Goranov *et al*., 2005). Increased dNTPs have been proposed to promote the RecA-dependent repair of stalled replication forks in *E. coli* (Robu, Inman, and Cox, 2001). This mechanism supports the model in which enhanced replication elongation helps overcome replication fork collapse caused by the severe overinitiation of Δ*yabA dnaA1*.

As mentioned before, changes in one step of DNA replication may affect another. For example, a small decrease in initiation could prevent replication fork collapse, indirectly allowing for increased elongation. This could be a possible mechanism for the *ccrZ* suppressors, where there is a decrease in *ori/ter* but no obvious change in DNA synthesis. This could also contribute to the mechanism of suppression by other mutants, however based on what is known about some genes identified, we suspect there are at least some direct effects to stimulate elongation or the repair of collapsed replication forks.

The diverse mutants isolated during this suppressor screen highlight the varied means that can be used to control detrimental DNA replication in a cell. This study emphasizes how the cell can act at different steps in DNA replication using multiple mechanisms to overcome a replication defect.

## Methods

### Media and growth conditions

Cells were grown with shaking at 37°C in Luria-Bertani (LB) medium (Miller, 1972) or S7 defined minimal medium with MOPS (3-(N-morpholino) propanesulfonic acid) buffer at a concentration of 50 mM rather than 100 mM supplemented with 1% glucose, 0.1% glutamate, trace metals, 40 µg/ml phenylalanine, and 40 ug/ml tryptophan (Jaacks *et al*., 1989). Standard concentrations of antibiotics were used when appropriate (Harwood and Cutting, 1990). To induce expression of *dnaC* from the LacI-repressible, IPTG-inducible Pspank, varied concentrations of IPTG were used as specified.

### Strains and alleles

The *E. coli* strain AG1111 (MC1061 F’ *lacI*^q^ *lacZ*M15 Tn*10*) (Glaser *et al*., 1993) was used for plasmid construction. *B. subtilis* strains were derived from JH642 (*pheA1 trpC2*) (Perego, Spiegelman, and Hoch, 1988; Smith, Goldberg, and Grossman, 2014), are listed in Table 3, and were constructed by natural transformation using genomic DNA.

AG1866 was constructed by transforming AG174 with KPL2 genomic DNA and selecting for mls (macrolide-lincosamide-streptogramin B) resistance and screening for temperature resistance at 50°C, resulting in a wild type strain isogenic to KPL2 (Tn917 Ω HU163 (mls)). The Tn917 Ω HU163 allele is inserted in *dcK* and linked to *dnaA*.

Δ*yabA*::*spc* (CAL2055) was constructed by replacing the *yabA* open reading frame in AG174 with a spectinomycin resistance cassette using long-flanking homology PCR. MEA64 is the result of backcrossing CAL2055 into AG1866 to generate a Δ*yabA*::*spc* strain isogenic to KPL2 and AG1866.

Deletions in *nrdR* (MEA187), *cshA* (MEA250), *purA* (MEA359) and *ytmP* (MEA587) were constructed by replacing the open reading frames with a kanamycin resistance cassette (*kan*) using linear Gibson isothermal assembly (Gibson *et al*., 2009) of fragments containing ∼1kb of flanking homology for the indicated gene.

P*dnaC*(*T7*) was reconstructed in a wild type background as follows: DNA from one of the originally isolated suppressors with the P*dnaC*(*T7*) mutation (MEA73) was transformed into a strain with Δ*purA*::*kan* (MEA359; linked to *dnaC* and an adenine auxotroph) selecting for adenine prototrophy on minimal medium plates. Candidates were sequenced to find isolates with the P*dnaC*(*T7*) mutation, resulting in strain MEA362.

CAL2320 was constructed by transforming CAL2055 into KPL2 and selecting on minimal medium plates with spectinomycin and confirming the presence of *dnaA1* by screening for t.s. at 50°C. All derivatives of CAL2320 (except MEA370) were constructed by introducing the desired alleles into KPL2 (selecting on rich medium) and then moving Δ*yabA*::*spc* (CAL2055) in last, selecting on minimal medium plates to avoid generating suppressors. MEA370 was constructed by moving *dnaA1*-Tn*917*ΩHU163 (*mls*) into MEA362 selecting on rich medium. Δ*yabA*::*spc* (CAL2055) was then moved in last, selecting on minimal medium.

MEA360 was constructed by transforming linearized pMEA358 (cut with DraIII) into AG174. pMEA358 is a derivative of pBOSE1404 that places the open reading frame (and ribosome binding site) of *dnaC* under the IPTG-inducible promoter Pspank. pBOSE1404 is a plasmid that introduces a chloramphenicol resistant Pspank at the *ycgO* locus (*ycgO*-*cat-lacI-*Pspank*-ycgO*) by double crossover. oMEA260 (5’-TTGTGAGCGGATAACAATTAAATGAGAGGACGGTGCTTAGC-3’) and oMEA261 (5’-GAATTAGCTTGCATGCGGATCTTTCGAATGAAAAAAACCCCAAGAG-3’) were used to amplify a 1420 bp fragment (AG174 as the template) that includes the open reading frame of *dnaC*. Using isothermal assembly, this fragment was introduced into pBOSE1404 cut with NheI and HindIII, generating pMEA358. MEA361 was constructed by replacing the open reading frame of *dnaC* with a kanamycin resistance cassette (kan) by using linear Gibson isothermal assembly fragments containing ∼1KB of flanking homology transformed into MEA360, selecting on kanamycin + 1mM IPTG to induce expression of the essential *dnaC*.

### Suppressor screen

Independent cultures of CAL2320 (Δ*yabA dnaA1*) were grown in defined minimal medium with 1% glucose (see above) at 37°C to mid exponential phase. Dilutions of each independent culture were plated on LB agar at 37°C. A single colony was chosen from each of the original independent cultures and streaked 2x on LB agar. Colonies with different growth rates and morphologies were chosen deliberately with the aim to diversify the mutants isolated.

### Genome sequencing

Each suppressor mutant and CAL2320 were grown in minimal medium with 1% glucose to mid exponential phase. Cells were harvested by centrifugation and DNA was isolated using a Qiagen 100 G tips purification kit. DNA was sheared using a Covaris ultrasonicator. Sample preparation, including incorporation of a 3’ barcode, selection of 300–600 bp fragments (after addition of adaptors and amplification), and paired-end read sequencing (150-150 nt) on an Illumina MiSeq were performed by the MIT BioMicro Center. Reads were mapped to the *B. subtilis* strain JH642 (GenBank: CP007800.1; Smith, Goldberg, and Grossman, 2014) as described (Deatherage and Barrick, 2014).

### qPCR to determine *ori/ter* ratio

Cultures were grown to mid-exponential phase and diluted back to OD 0.05 and grown to mid-exponential phase (OD 0.2-0.4) in defined minimal medium at 37°C. Cells were harvested in ice-cold methanol (1:1 ratio) and pelleted. Genomic DNA was isolated using Qiagen DNeasy kit with 40 µg/ml lysozyme. The copy number of the origin (*ori*) and terminus (*ter*) were quantified by qPCR to generate an *ori/ter* ratio. qPCR was done using SSoAdvanced SYBR master mix and CFX96 Touch Real-Time PCR system (Bio-Rad). Primers used to quantify the origin region were oMEA316 (5’-TTGCCGCAGATTGAAGAG-3’) and oMEA317 (5’-AGGTGGACACTGCAAATAC-3’). Primers used to quantify the terminus region were oMEA318 (5’-CGCGCTGACTCTGATATTATG-3’) and oMEA319 (5’-CAAAGAGGAGCTGCTGTAAC-3’). Origin-to-terminus ratios were determined by dividing the number of copies (as indicated by the Cp values measured through qPCRs) of the origin by the number of copies quantified at the terminus. Ratios were normalized to the origin-to-terminus ratio of a temperature sensitive mutant, *dnaB134* (KPL69), that was shifted to the non-permissive temperature, which allows ongoing replication to be completed but does not allow new initiation, resulting in 1:1 ratio of the origin:terminus.

### RT-qPCR

Cultures were grown to mid-exponential phase and diluted back to OD 0.05 and grown to mid-exponential phase (OD 0.2-0.4) in defined minimal medium at 37°C. Cells were harvested in ice-cold methanol (1:1 ratio) and pelleted. RNA was isolated using Qiagen RNeasy PLUS kit with 10 mg/ml lysozyme. iScript Supermix (Bio-Rad) was used for reverse transcriptase reactions to generate cDNA. RNA was degraded by adding 75% volume of 0.1 M NaOH and incubating at 70°C 10 minutes, followed by neutralizing the reaction with adding 75% of the original volume 0.1 M HCl. qPCR was done using SSoAdvanced SYBR master mix and CFX96 Touch Real-Time PCR system (Bio-Rad). Primers used to quantify *dnaC* were oMEA126 (5’-AGCTGCAAGTCCCTGTTATC-3’) and oMEA127 (5’-CCTGCTCGATACTTCCTGATTC-3’). Primers used to quantify *dinC* were oMEA207 (5’-ACCAAGATACACCTCCAGAAAG-3’) and oMEA208 (5’-AACCTGAAGTCGGAACCATC-3’). Primers used to quantify *sigA* were oMEA252 (5’-ATACCGGCTCTTGAGCAATC-3’) and oMEA253 (5’-ACTTAGGCAGAGAACCAACAC-3’). Primers used to quantify *gryA* were oMEA128 (5’-TGGAGCATTACCTTGACCATC-3’) and oMEA129 (5’-AGCTCTCGCTTCTGCTTTAC-3’).

### Western blots to measure protein levels

Cultures were grown to mid-exponential phase and diluted back to OD 0.05 and grown to mid-exponential phase (OD 0.2-0.4) in defined minimal medium or LB (as indicated) at 37°C. Exponentially growing cells were lysed with lysozyme and a protease inhibitor cocktail (Sigma-Aldrich, P8849) at 37°C. Lysates were run on 12% polyacrylamide gel and transferred to a nitrocellulose membrane using the Trans-blot SD semi-dry transfer cell (Biorad). To represent the same number of cells in each lane, the amounts loaded in each lane were normalized to culture OD. All steps were performed at room temperature. According to the manufacturer’s instructions, blots were blocked with Odyssey Blocking Buffer for 1 hr and incubated with primary antibody (rabbit anti-DnaC antibody 1:10,000 in Odyssey Blocking Buffer + 0.2% Tween) for 1 hr. The primary antibody to DnaC was affinity purified rabbit polyclonal antibody (Covance) made against purified DnaC-His6. Blots were washed with PBST (phosphate-buffered saline + 0.2% Tween) for at least 5 x 5 min and then incubated for 30 min with secondary antibody (LiCor dye 800 goat anti-rabbit 1:10,000 in Odyssey Blocking Buffer + 0.2% Tween). Blots were imaged and quantitated on a LiCor scanner. Dilutions of AG174 lysates were used to generate a standard curve and determine the linear range of the fluorescence signal.

### ^3^H-thymidine incorporation to measure DNA synthesis

Cells were grown at 37°C, and were assayed at 3 to 5 time-points between OD 0.2 and 0.5. Incorporation per OD was found to be linear over this range. A 250 µl aliquot was pulse labeled with 10 µl 3H-thymidine (6.7 Ci/mmol; 1 mCi/ml). After 1 min, a short chase was performed by adding 50 µl 10 mM unlabeled thymidine and incubating for an additional 1 min. An equal volume ice-cold 20% trichloroacetic acid (TCA) was added, and the samples were incubated on ice for 30-60 min. A 350 µl aliquot was vacuum-filtered on glass-fiber filters (24 mm GF/A, Whatman) and washed with 25 ml of ice-cold 5% TCA, followed by 2 ml 100% ethanol. The amount of radioactivity that had been incorporated into nucleic acid was determined by scintillation counting of the dried filters. A time course was performed to confirm that the 1 min pulse was in the linear range of the assay.

### Mass spectrometry to measure nucleotide pools

Cultures were grown in minimal glucose media to OD 0.45-0.55, transferred to 50 ml conical tube, and quickly chilled to 10°C by swirling in liquid nitrogen. Cells from 5 ml of the chilled culture were collected onto a 25 mm MilliPore Type HAWP 0.45 µM filter using a filtration apparatus chilled to 4°C. The filter was immediately placed in a tube containing 1 ml ice-cold extraction buffer (40% methanol, 40% acetonitrile, 20% water with 0.1M formic acid, and 500 nM each of 17 isotopically-labeled amino acids), and quickly frozen in liquid nitrogen. After removing from liquid nitrogen, 87µl 15% ammonium bicarbonate was added to each tube, and vortexed and shaken vigorously to disrupt the cells. After 5 min centrifugation at 16100 g at 4°C, the supernatant was removed and dried under vacuum. LC-MS profiling and analysis was performed by the Whitehead Institute Proteomics Core Facility essentially as described (Kanarek *et al*., 2018). Raw LC-MS data were analyzed using MZMine 2 (Pluskal, *et al*., 2010). dUMP and dTMP peak areas were corrected for small differences in the ODs of cells processed, and normalized to the glutamate internal standard.

## Acknowledgements

We thank Caroline Lewis, Tenzin Kunchok, and Bena Chan at the Metabolite Profiling Core Facility at the Whitehead Institute for running metabolomics samples and for help with data analysis. We thank Stuart Levine and the BioMicro Center at MIT for help with the genome sequencing. We thank Stephen Bell, Michael Laub, and Gene-Wei Li for helpful discussions and comments on the manuscript.

Research reported here is based upon work supported, in part, by the National Institute of General Medical Sciences of the National Institutes of Health under award number R01 GM041934 and R35 GM122538 to ADG. Any opinions, findings, and conclusions or recommendations expressed in this report are those of the authors and do not necessarily reflect the views of the National Institutes of Health.

